# Astrocyte Ca^2+^ Signaling is Facilitated in an *Scn1a^+/−^* Mouse Model of Dravet Syndrome

**DOI:** 10.1101/2021.05.18.444602

**Authors:** Kouya Uchino, Wakana Ikezawa, Yasuyoshi Tanaka, Masanobu Deshimaru, Kaori Kubota, Takuya Watanabe, Shutaro Katsurabayashi, Katsunori Iwasaki, Shinichi Hirose

## Abstract

Dravet syndrome (DS) is an infantile-onset epileptic encephalopathy. More than 80% of DS patients have a heterozygous mutation in *SCN1A*, which encodes a subunit of the voltage-gated sodium channel, Nav_1.1_, in neurons. The roles played by astrocytes, the most abundant glial cell type in the brain, have been investigated in the pathogenesis of epilepsy; however, the specific involvement of astrocytes in DS has not been clarified. In this study, we evaluated Ca^2+^ signaling in astrocytes using genetically modified mice that have a loss-of-function mutation in *Scn1a*. We found that the slope of spontaneous Ca^2+^ spiking was increased without a change in amplitude in *Scn1a^+/−^* astrocytes. In addition, ATP-induced transient Ca^2+^ influx and the slope of Ca^2+^ spiking were also increased in *Scn1a^+/−^* astrocytes. These data indicate that perturbed Ca^2+^ dynamics in astrocytes may be involved in the pathogenesis of DS.

## Introduction

Epilepsy is a chronic neurological disorder that causes paroxysmal loss of consciousness and convulsions because of overexcitement of neurons in the brain (1). Antiepileptic drugs are commonly used to treat epilepsy, but 20%–30% of patients show resistance to drug therapy (2). Dravet syndrome (DS) is an epileptic encephalopathy that occurs in infancy at a frequency of 1 in 20,000 to 40,000 children (3, 4). In addition, 70%–80% of DS patients have heterozygous mutations in sodium voltage-gated channel alpha subunit 1 (*SCN1A*), which encodes a subunit of the voltage-gated sodium channel, Nav_1.1_ (5). A mouse model with a heterozygous loss-of-function mutation in *Scn1a* was recently reported as a DS model (6–9). We recently reported that the excitatory and inhibitory balance of synaptic transmission was disrupted in heterozygous *Scn1a* knockin (*Scn1a^+/−^*) mouse neurons when the extracellular Ca^2+^ concentration was increased (10). The above studies show that mutations in the *Scn1a* gene cause abnormal neurological function. In addition to neuronal dysfunction, the involvement of glial cells in the pathogenesis of epilepsy has also been suggested (11). In particular, astrocytes, a type of glial cell, are involved in epilepsy pathogenesis by regulating neurotransmitter and ion concentrations (12). Therefore, astrocytes are a promising new therapeutic target for epilepsy.

Unlike neurons, which transmit information by generating action potentials, astrocytes do not generate action potentials and have been considered non-excitable cells. However, astrocytes are found throughout the central nervous system and play an essential role in neuronal function (13). For example, we have demonstrated that the long-term culture of astrocytes and changes in astrocyte density can alter neuronal growth and synaptic transmission (14, 15). Advances in Ca^2+^ imaging methods have revealed that astrocytes are excitable cells that exhibit intracellular Ca^2+^ signaling (16). Astrocytes regulate neuronal activity by releasing glial transmitters and neurotransmitters, such as glutamate, ATP, and D-serine (17). In addition, astrocyte Ca^2+^ signaling is involved in epileptic seizures via neurons (18, 19). *Scn1a* mutations change the neuronal expression of Ca^2+^ channels and sensitivity to Ca^2+^ (10, 20); however, it is not known if astrocytes are affected in the DS model. In this study, we evaluated Ca^2+^ dynamics in astrocytes using genetically modified *Scn1a^+/−^* mice.

## Materials and Methods

### Animals

Experimental animals were handled in accordance with the ethical regulations for animal experiments of the Fukuoka University Experimental Animal Care and Use Committee. All animal protocols were approved by the Ethics Committee of Fukuoka University (permit numbers: 1712128 and 1812092). All experimental protocols were performed according to the relevant guidelines and regulations of Fukuoka University. All *in vivo* work was carried out in compliance with ARRIVE guidelines. *Scn1a^+/−^* mice, in which coding exons 8–12 of *Scn1a* were replaced with a neomycin resistance gene, were generated as previously described (10). The mice were housed in plastic cages and kept at 23±2°C, in 60±2% humidity, and with a 12-hour light/dark cycle (lights on at 7:00 am, lights off 7:00 pm). Food (CE-2, CLEA Japan, Inc., Tokyo, Japan) and water were freely available.

### Astrocyte culture

For astrocyte culture, newborn *Scn1a^+/−^* mice (P0–1) were used. Tail biopsies were taken and PCR performed to confirm the *Scn1a* genotype (10). Cerebral cortices of 0–1-day-old mice were treated with trypsin/EDTA (0.05%/0.02%) and cultured in D-MEM medium containing 10% fetal bovine serum at 37°C for 2 weeks. The astrocyte layer adhering to the bottom of the culture bottle was then detached with trypsin/EDTA (0.05%/0.02%). Cultured astrocytes were seeded at a density of 75,000 cells/well in six-well plates on glass coverslips coated with collagen/poly-D-lysine and cultured for 7–8 days.

### Ca^2+^ imaging

Spontaneous Ca^2+^ signaling was monitored using Oregon-Green BAPTA-1 AM, a fluorescent Ca^2+^ indicator (Thermo Fisher Scientific, Waltham, MA, USA). Oregon-Green BAPTA-1 AM emits green fluorescence at resting Ca^2+^ levels, and the fluorescence intensity increases with increasing Ca^2+^ binding. ATP-induced Ca^2+^ signaling was monitored using Fluo4-AM, a fluorescent Ca^2+^ indicator that does not have a resting signal (Thermo Fisher Scientific). Fluo4-AM shows minimal fluorescence at resting Ca^2+^ levels, and the fluorescence emission intensity increases with Ca^2+^ binding.

Astrocytes were incubated with Oregon-Green BAPTA-1, AM, or Fluo4-AM for 60 min. Astrocytes were incubated with NucBlue (Thermo Fisher Scientific) for 20 min to identify nuclei. The glass coverslips were then transferred to a recording chamber and refluxed with an extracellular solution. Fluorescence excitation was performed using LED (light-emitting diode) irradiation (Lambda HPX, Sutter Instrument, Novato, CA, USA). For time-lapse imaging, the LED flash and the exposure time of an sCMOS camera (edge4.2, pco, Kelheim, Germany) were synchronized. Oregon-Green BAPTA-1, AM was irradiated with 494 nm excitation light and observed with a fluorescence wavelength of 523 nm (exposure time 100 ms, interval 1000 ms, number of images 360). Fluo4-AM was irradiated with 495 nm excitation light and observed with a fluorescence wavelength of 518 nm (exposure time 100 ms, interval 1000 ms, number of images 360). For Fluo4-AM observation, 1 μM ATP was applied for 30 s to induce Ca^2+^ signaling.

### Data analysis

ImageJ software (1.53c, Wayne Rasband, NIH, USA) and AxoGraph X software (1.2, AxoGraph Scientific, Sydney, Australia) were used to analyze the dynamics of Ca^2+^ signaling. Images stained with NucBlue were divided into nuclei and background by threshold; a unit of >100 pixels was considered the location of a nucleus and registered as a region of interest (ROI). The ROIs were then fitted to the stacked images, and the intensity change was measured. The maximum intensity value in the stack image for each nucleus was transferred to Axograph. For analysis of spontaneous Ca^2+^ signaling, the data value was relativized in Axograph, and the maximum relative intensity was calculated. The data were differentiated from the relative value, and the maximum differential intensity was obtained. For analysis of ATP-induced Ca^2+^ signaling, the intensity (F) of each ROI was normalized by the intensity before ATP application (F0). The area under the curve (AUC) of intensity was also measured from the ATP-induced Ca^2+^ wave. In addition, the F/F0 was differentiated, and the maximum differential intensity was calculated.

### Solutions

Extracellular solution (pH 7.4) consisted of (mM): NaCl 140, KCl 2.4, HEPES 10, glucose 10, CaCl_2_ 2, and MgCl_2_ 1. ATP (Thermo Fisher Scientific) was dissolved in ultrapure water to make a 100-mM stock solution. The stock was diluted in extracellular solution to 1 μM just before use.

### Statistics

All data are expressed as the mean ± standard error. A lowercase n indicates the number of astrocytes recorded, and an uppercase N indicates the number of cultures (lot number). Two groups were compared with Student’s unpaired t-test using Kaleida Graph 3.6 (Synergy Software, Reading, PA, USA). Statistical significance was considered when p < 0.05.

## Results

Astrocytes regulate neurotransmitter and ion concentrations in neurons and other glial cells via Ca^2+^ signaling. The concentrations of many neurotransmitters and ions are abnormal in epilepsy; therefore, we expected that Ca^2+^ signaling would be altered in *Scn1a^+/−^* astrocytes. Oregon-Green BAPTA-1, AM detects the resting signal of Ca^2+^, meaning that it was not possible to fix a baseline or to measure the frequency of spontaneous Ca^2+^ spiking because of baseline instability. Therefore, we first obtained the relative strength of spontaneous Ca^2+^ signaling in cultured astrocytes from the cerebral cortex of *Scn1a^+/−^* mice. The maximum relative intensity of spontaneous Ca^2+^ signaling was identical between wild-type (WT) and *Scn1a^+/−^* astrocytes (Figures 1A, B, C). To analyze the Ca^2+^ waveform in more detail, we differentiated the waveform of spontaneous Ca^2+^ signaling. As a result, the maximum differential intensity was significantly increased in *Scn1a^+/−^* compared with WT astrocytes (Figures 1D, E). These results indicate that the unitary speed of Ca^2+^ spiking was increased without change in the amount of spontaneous Ca^2+^ influx in *Scn1a^+/−^* astrocytes.

**Figure 1.**
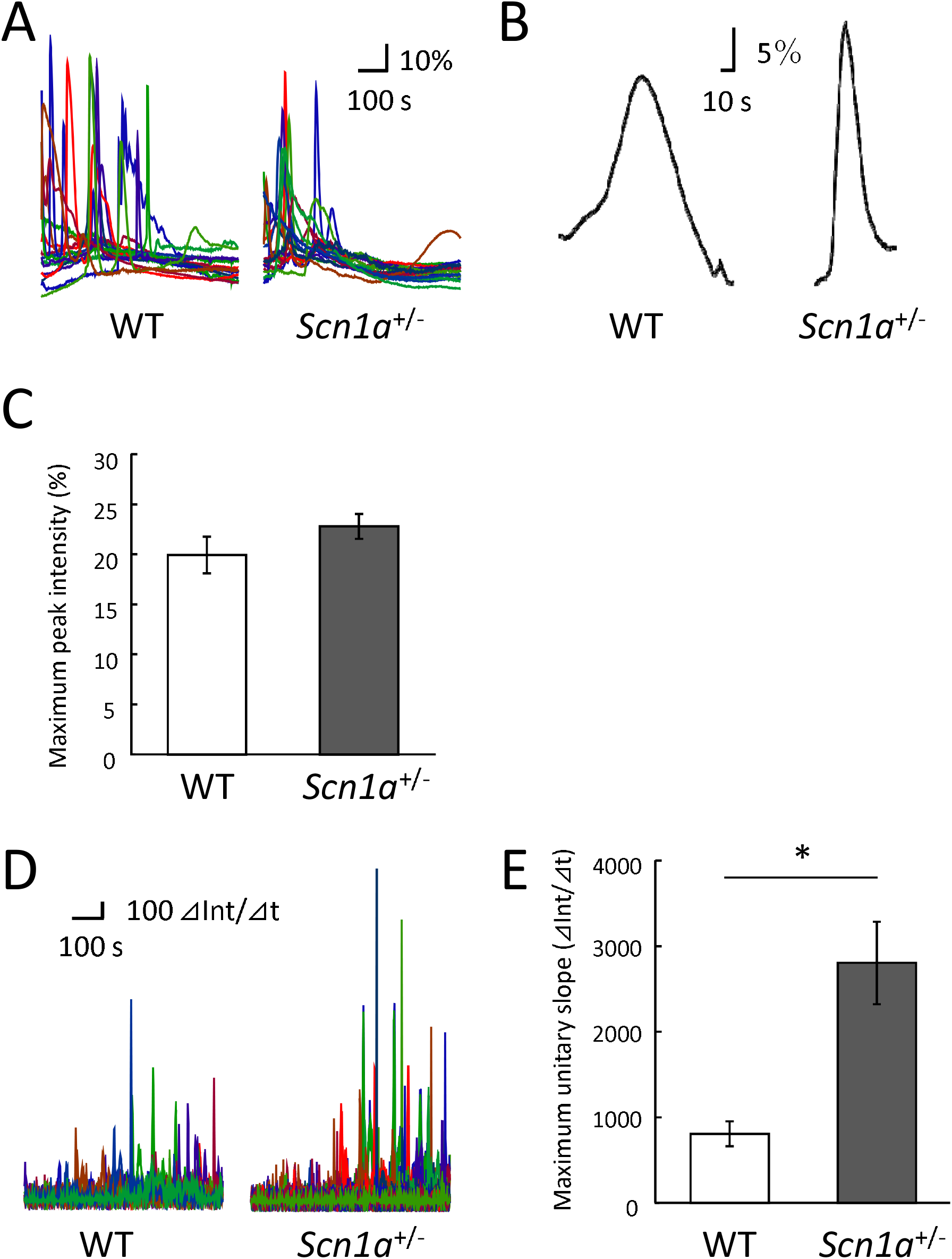
Spontaneous Ca^2+^ signaling in *Scn1a^+/−^* astrocytes. (A) Representative waveforms of spontaneous Ca^2+^ signaling in wild-type (WT) and *Scn1a^+/−^* astrocytes. Waveforms are shown relative to the baseline. (B) The waveform expanded from a part of (A). (C) Mean of maximum relative intensity of spontaneous Ca^2+^ signaling in WT and *Scn1a^+/−^* astrocytes (WT: 19.9 ± 1.8%, n = 123; *Scn1a^+/−^*: 22.8 ± 1.3%, n = 179 from N = 5 cultures). (D) Representative differentiated waveforms of spontaneous Ca^2+^ signaling in WT and *Scn1a^+/−^* astrocytes. (E) Mean of the maximum differential intensity of spontaneous Ca^2+^ signaling in WT and *Scn1a^+/−^* astrocytes (WT: 797.5 ± 143.2 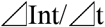, n = 98; *Scn1a^+/−^*: 2796 ± 481.3 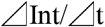, n = 179 from N = 4 cultures).

In epilepsy, external stimuli may trigger neuronal hyperexcitability, resulting in epileptic seizures. Therefore, we stimulated astrocytes with ATP (1 μM) and recorded the ATP-induced Ca^2+^ signaling. As shown in Figure 2, the maximum peak intensity was dramatically increased in *Scn1a^+/−^* astrocytes (Figures 2A, B, C). However, there was no significant difference in the AUC of ATP-induced Ca^2+^ signaling between WT and *Scn1a^+/−^* astrocytes (Figure 2D). These data indicate that *Scn1a^+/−^* astrocytes exhibit a sharper Ca^2+^ waveform. We then analyzed the slope of the wave by differentiating the Ca^2+^ waveform and, as expected, the maximum differential intensity significantly increased in *Scn1a^+/−^* astrocytes (Figures 2E, F). These results indicate that the transient dynamics of Ca^2+^ spiking by ATP stimulation were facilitated without changing the total amount of Ca^2+^ signaling in *Scn1a^+/−^* astrocytes.

**Figure 2.**
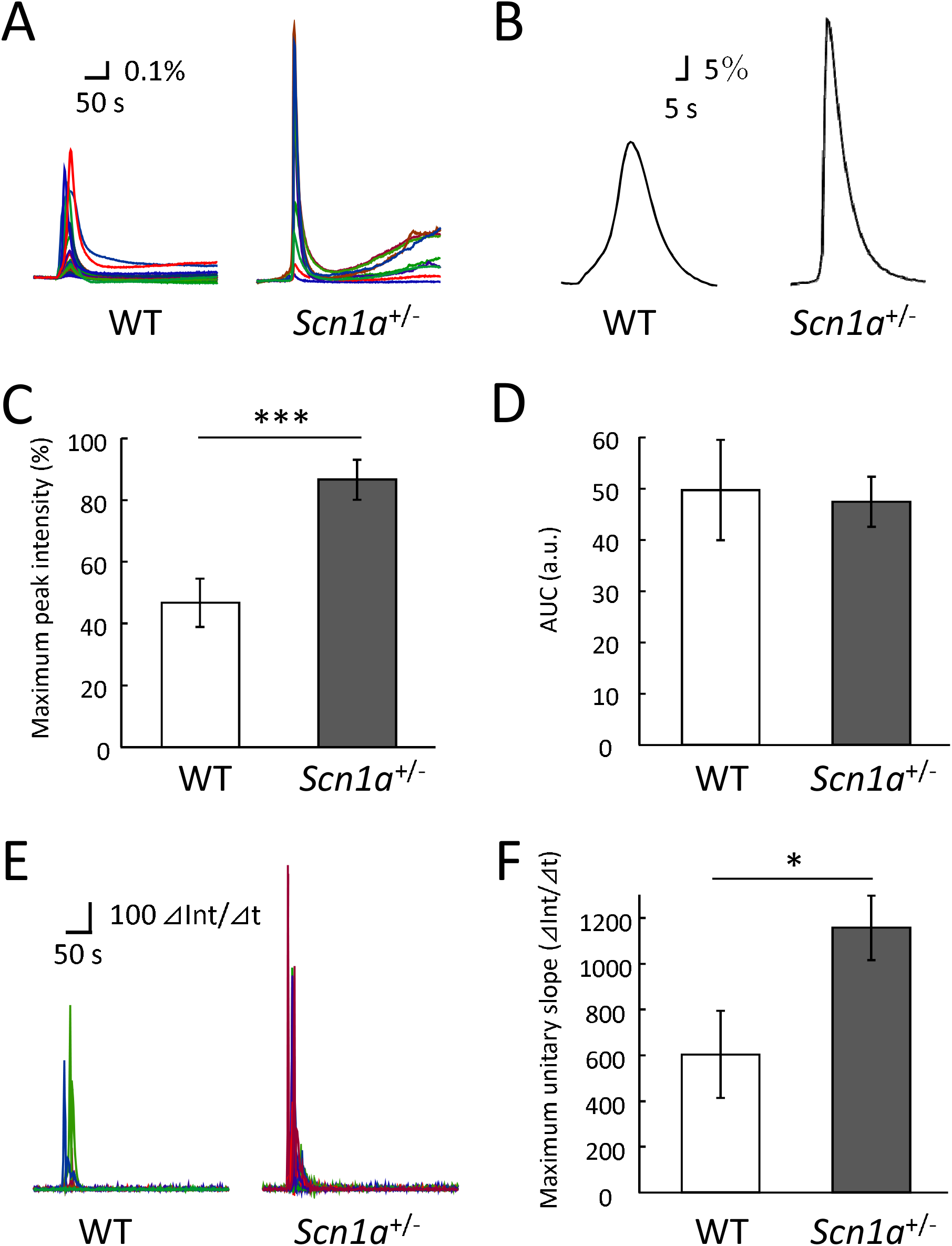
ATP-induced Ca^2+^ signaling in *Scn1a^+/−^* astrocytes. (A) Representative waveforms of ATP-induced Ca^2+^ signaling in WT and *Scn1a^+/−^* astrocytes. Waveforms are shown relative to F/F0. (B) The waveform expanded from a part of (A). (C) Mean of maximum relative peak intensity of ATP-induced Ca^2+^ signaling in WT and *Scn1a^+/−^* astrocytes (WT: 46.5 ± 7.8%, n = 64; *Scn1a^+/−^*: 86.4 ± 6.5%, n = 114 from N = 4 cultures). (D) Mean AUC of ATP-induced Ca^2+^ signaling in WT and *Scn1a^+/−^* astrocytes (WT: 49.5 ± 9.8 a.u., n = 74; *Scn1a^+/−^*: 47.3 ± 4.9 a.u., n = 125 from N = 4 cultures). (E) Representative differentiated waveforms of ATP-induced Ca^2+^ signaling in WT and *Scn1a^+/−^* astrocytes. (F) Mean values of the maximum differential intensity of ATP-induced Ca^2+^ signaling in WT and *Scn1a^+/−^* astrocytes (WT: 601.1 ± 190.2 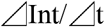, n = 63; *Scn1a^+/−^*: 1154.5 ± 140.7 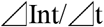, n = 114 from N = 4 cultures).

## Discussion

Astrocytes regulate the concentrations of neurotransmitters and ions in neurons and other glial cells via Ca^2+^ signaling. The concentrations of many neurotransmitters and ions are abnormal in epilepsy (21); therefore, we expected that Ca^2+^ signaling in astrocytes would be disrupted in the DS mouse model. Our results show that the transient spontaneous influx of Ca^2+^ did not change, but that the slope of Ca^2+^ spiking increased in *Scn1a^+/−^* astrocytes. Likewise, in response to ATP stimulation, there was no change in the total amount of Ca^2+^ signaling, but the transient Ca^2+^ influx and the slope of Ca^2+^ spiking increased in *Scn1a^+/−^* astrocytes. These results indicate that Ca^2+^ signaling is somehow enhanced in *Scn1a* astrocytes. Spontaneous Ca^2+^ signaling in astrocytes is thought to result from the uptake of extracellular Ca^2+^. In contrast, stimulus-induced Ca^2+^ signaling is thought to be caused by the release of Ca^2+^ into the endoplasmic reticulum via the activation of Gq protein-coupled receptors (17). In the present study, both spontaneous and ATP-induced Ca^2+^ signaling were enhanced in *Scn1a^+/−^* astrocytes, indicating that the function of both cascades may be enhanced.

Since it was first reported that the concentration of cytoplasmic Ca^2+^ in cultured astrocytes increases in response to glutamate (22), the concept of electrically unresponsive astrocytes sensing glutamate transmission has been accepted. This observation, and that the increase in astrocyte Ca^2+^ is associated with neurotransmitter release, is the basis for the concept of tripartite synapses (23–25). Astrocytes release glutamate via metabotropic glutamate receptors (mGluRs), which is thought to regulate the amount of glutamate in the brain, although this is a controversial assumption (26, 27). The expression of mGluRs in astrocytes is increased in mouse models of epilepsy (28), indicating that mGluR activation contributes to epileptic seizures caused by excess glutamate release in the brain (29). However, ATP released upon Ca^2+^ signaling is degraded to adenosine; therefore, adenosine may suppress epileptic seizures by exerting an inhibitory effect on neurons through pre-synaptic A1 receptors (30).

In conclusion, it is not known whether enhanced astrocyte Ca^2+^ signaling exacerbates epileptic seizures or suppresses them. However, the present study indicates that astrocytes are involved in the pathogenesis of DS. This may lead to novel pharmaceutical treatments that target non-traditional mechanisms. We suggest that study of the interaction between astrocytes and neurons, focusing on the clinical context in humans, is warranted.

## Contribution to the field

Dravet syndrome (DS) usually begins in the first year of life and is a severe form of epilepsy that often leads to severe encephalopathy. Over 80% of DS patients have a heterozygous mutation in *SCN1A* (which encodes a subunit of voltage-gated Na^+^ channels). However, the mechanisms underlying this disease remain unknown, which makes drug development difficult.

In addition to neuronal involvement, astrocytes—the most abundant glial cell type in the brain—are also involved in the pathogenesis of epilepsy. Therefore, astrocytes are attracting attention as a new therapeutic target for epilepsy. In this study, we found that Ca^2+^ spiking was significantly faster and that ATP-induced Ca^2+^ spiking was more significant in astrocytes cultured from *Scn1a^+/−^* mice compared with that in astrocytes from wild-type mice.

This paper demonstrates that Ca^2+^ dynamics in astrocytes may be involved in the pathogenesis of DS. Astrocytes play a vital role in protecting neural circuits; therefore, the changes we identified in *Scn1a^+/−^* astrocyte Ca^2+^ signaling may help in the development of novel therapies for epilepsy that target astrocytes to protect neural circuits.

## Acknowledgements

We thank the members of our laboratory for their assistance. We thank Jeremy Allen, PhD, from Edanz (https://jp.edanz.com/ac) for editing a draft of this manuscript.

## Data Availability Statement

The data included in this study are available from the corresponding author on reasonable request.

## Ethics Statement

We confirm that we have read the Journal’s position on issues involved in ethical publication and affirm that this report is consistent with these guidelines.

## Author Contributions

K.U., W.I. and Y.T. performed experiments and analyzed data; Y.T. and M.D. created the *Scn1a^+/−^* mouse model; S.K. conceived the study; K.K., T.W., K.I., and S.H. interpreted the data; K.U. and S.K. wrote the manuscript with input from all authors. All authors reviewed the manuscript.

